# On the causes of evolutionary transition:transversion bias

**DOI:** 10.1101/027722

**Authors:** Arlin Stoltzfus, Ryan W. Norris

## Abstract

A pattern in which nucleotide transitions are favored several-fold over transversions is common in molecular evolution. When this pattern occurs among amino acid replacements, explanations often invoke an effect of selection, on the grounds that transitions are more conservative in their effects on proteins. However, the underlying hypothesis of conservative transitions has never been tested directly. Here we assess support for this hypothesis using direct evidence: the fitness effects of mutations in actual proteins, measured via individual or paired growth experiments. We assembled data from 8 published studies, ranging in size from 24 to 757 single-nucleotide mutations that change an amino acid. Every study has the statistical power to reveal significant effects of amino acid exchangeability, and most studies have the power to discern a binary conservative-vs-radical distinction. However, only one study suggests that transitions are significantly more conservative than transversions. In the combined set of 1239 replacements, the chance that a transition is more conservative than a transversion is 53 % (95 % confidence interval, 50 % to 56 %), compared to the null expectation of 50 %. We show that this effect is not large compared to that of most biochemical factors, and is not large enough to explain the several-fold bias observed in evolution. In short, available data have the power to verify the “conservative transitions” hypothesis if true, but suggest instead that selection on proteins plays at best a minor role in the observed bias.

## Introduction

Of the 12 types of changes from one nucleotide to another, 8 are “transversions” between a purine (A or G) and a pyrimidine (C or T), and the other 4 are “transitions”. Early protein comparisons showed that related proteins often differ by transitions more than expected by chance (e.g., Fitch 1967; sources cited in Vogel 1972). By the 1980's, this “transition bias” was well known (Li, et al. 1985). By the 1990s, systematists had noted effects on phylogeny inference (Wakeley 1996), and methods were revised to give more weight to transversion differences (e.g., Sinsheimer, et al. 1997).

In many early works, this bias is presented as a ratio of differences, which makes the expected ratio a complex function of the degree of sequence divergence. As the use of rate models became routine in comparative sequence analysis, the phenomenon of transition bias was redefined as a bias in instantaneous rates, relative to a null model of equal rates. Because every nucleotide site (e.g., a G site) may experience 1 type of transition (G→A) at rate α, and 2 types of transversion (G→C, G→T) at rate β, the aggregate rate ratio of transitions to transversions has a null expectation of R = α/(2β) = 0.5. In some contexts, the ratio is expressed differently as κ = α/β = 1. When considering amino acid changes, it is more relevant to compare the 116 possible transitions and 276 possible transversions that change a codon so as to encode a different amino acid (assuming the canonical genetic code), leading to a null expectation of R = 116 α / (276 β) = 0.42 α/β. Thus, the observation of roughly equal numbers of inferred transitions and transversions in classic works (e.g., Vogel and Kopun 1977), or in the extensive analysis of mammalian genes in Li (1997, Table 7.2), indicates a bias of over 2-fold. Kumar (1996) estimates 2-fold to 5-fold rate biases in vertebrate mitochondrial genes (excluding 3^rd^ positions). Other estimates may be found in work cited by Rosenberg, et al. (2003), but there is not (to our knowledge) a systematic contemporary review of this issue.

The causes of the observed bias have not been resolved. The hypothesis of a mutational cause— a transition:transversion bias in mutation— was promoted early by Vogel (1972; see also Vogel and Kopun 1977). This hypothesis was bolstered when DNA sequence comparisons revealed that a transition bias is observed in introns and other non-coding regions (Li, et al. 1985), suggesting a cause that (like mutation) acts at the level of DNA, across the entire genome.

The alternative hypothesis that natural selection favors amino acid replacements via transitions is also common, and is argued on the grounds that transitions are “less severe with respect to the chemical properties of the original and mutant amino acids” (Rosenberg, et al. 2003) or “tend to cause changes that conserve the chemical properties of amino acids” (Wakeley 1996), or that “the biochemical difference in the protein product tends to be greater for transversions” (Keller, et al. 2007).

For purposes of evaluation, we can break down either the mutational hypothesis or the selective hypothesis into (1) a claim that there is an underlying bias (mutational or selective) favoring transitions, and (2) a claim that this bias accounts for the observed evolutionary bias. For the mutational hypothesis, the existence of an underlying bias is indicated in direct studies of mutation (e.g., Schaaper and Dunn 1991; Lynch 2010; Schrider, et al. 2013; Zhu, et al. 2014), and by many indirect estimates based on the asumption of neutral sequence divergence (Petrov and Hartl 1999; Rosenberg, et al. 2003; Zhao, et al. 2004; Jiang and Zhao 2006; Morton, et al. 2006), though Keller, et al. (2007) report a lack of bias in grasshoppers. The bias typically is 2-fold to 4-fold over null expectations. In theory, a bias in mutation of magnitude B can cause a B-fold effect on the rate of evolution (Yampolsky and Stoltzfus 2001; McCandlish and Stoltzfus 2014). That is, the observed magnitude of mutation bias appears to be sufficient, in principle, to account for the observed evolutionary bias.

For the selective hypothesis, arguments to the effect that transitions are more conservative typically invoke a biochemical factor (or a composite such as the Grantham index) that correlates with patterns of evolutionary divergence, and is found to be more conserved by transitions than by transversions (e.g., Vogel and Kopun 1977; Zhang 2000). This form of argument suffers from a logical circularity: if mutation shapes patterns of evolutionary amino acid replacement, then biochemical factors chosen for their ability to make sense of evolutionary patterns are not independent of mutation.

Presumably no biochemical factor, nor any simple combination of factors, fully captures the effects of replacements in complex proteins operating in a complex milieu. Indeed, the use of biochemical surrogates would seem unnecessary, given the availability of more direct measurements. Systematic laboratory studies of the effects of amino acid replacements in proteins have been carried out for 25 years (e.g., Kleina and Miller 1990). Whereas early studies summarized by Yampolsky and Stoltzfus (2005) typically reported crude measures of biochemical or growth effects (e.g., a 2-valued scale of “−” and “+”), a number of more recent studies report a continuous measure of fitness for each mutant (e.g., Sanjuan, et al. 2004; Carrasco, et al. 2007; Domingo-Calap, et al. 2009; Peris, et al. 2010; Jacquier, et al. 2013; Roscoe, et al. 2013; Acevedo, et al. 2014; Bloom 2014; Firnberg, et al. 2014; Thyagarajan and Bloom 2014; Wu, et al. 2014). Such studies provide direct evidence on the relative conservativeness of transitions and transversions that change amino acids.

Here we focus on whether direct measurements of fitness support the conservative transitions hypothesis, based on a collection of 8 studies comprising measured fitness values for 544 transitions and 695 transversions that change an amino acid. We assess the power of each study by comparing mutant fitnesses for each type of replacement (e.g., Ser to Pro) with a cross-validation predictor and with 2 existing measures of amino acid exchangeability called EX (Yampolsky and Stoltzfus 2005) and U (Tang, et al. 2004). We find that, for every mutation study, even the smallest, there is a significant correlation with one or more of these predictors; half of the studies show a highly significant correlation (P < 0.001). More importantly, for most studies, measured fitness values correlate significantly with a conservative-vs-radical distinction based on EX or U. Specifically, a replacement designated as “conservative” has a 65 % (EX) or 64 % (U) chance of being more fit than a “radical” replacement.

However, the same studies typically do **not** show significant conservativeness of transitions. In the combined data, a transition has a 53 % chance (CI, 50 % to 56 %) of being more fit than a transversion, only slightly above the null expectation of 50 %. We show that this effect is not large compared to that of most biochemical predictors, and is not large enough to explain the several-fold bias toward transition replacements observed in evolutionary studies. The mutation-bias hypothesis, though not proven, remains an obvious possibility, while the selective hypothesis would seem untenable.

### Results

The literature search described in Materials and Methods (see also Supplementary Material) resulted in the 8 data sets in Table 1, each of which provides measures of fitness based on individual growth or paired growth (Sanjuan, et al. 2004; Carrasco, et al. 2007; Domingo-Calap, et al. 2009; Peris, et al. 2010; Jacquier, et al. 2013; Rihn, et al. 2013; Rihn, et al. 2015). We will refer to these 8 data sets as 8 studies, although they correspond to 7 publications, one of which (Domingo-Calap, et al. 2009) reports separate mutant fitness distributions for 2 different phages. Because measures of growth from different studies are not scaled in the same way, we convert fitnesses to within-study quantiles, e.g., the median fitness in a study is assigned a quantile of 0.5, and the fitness at the 95^th^ percentile is assigned a quantile of 0.95.

**Table 1.**
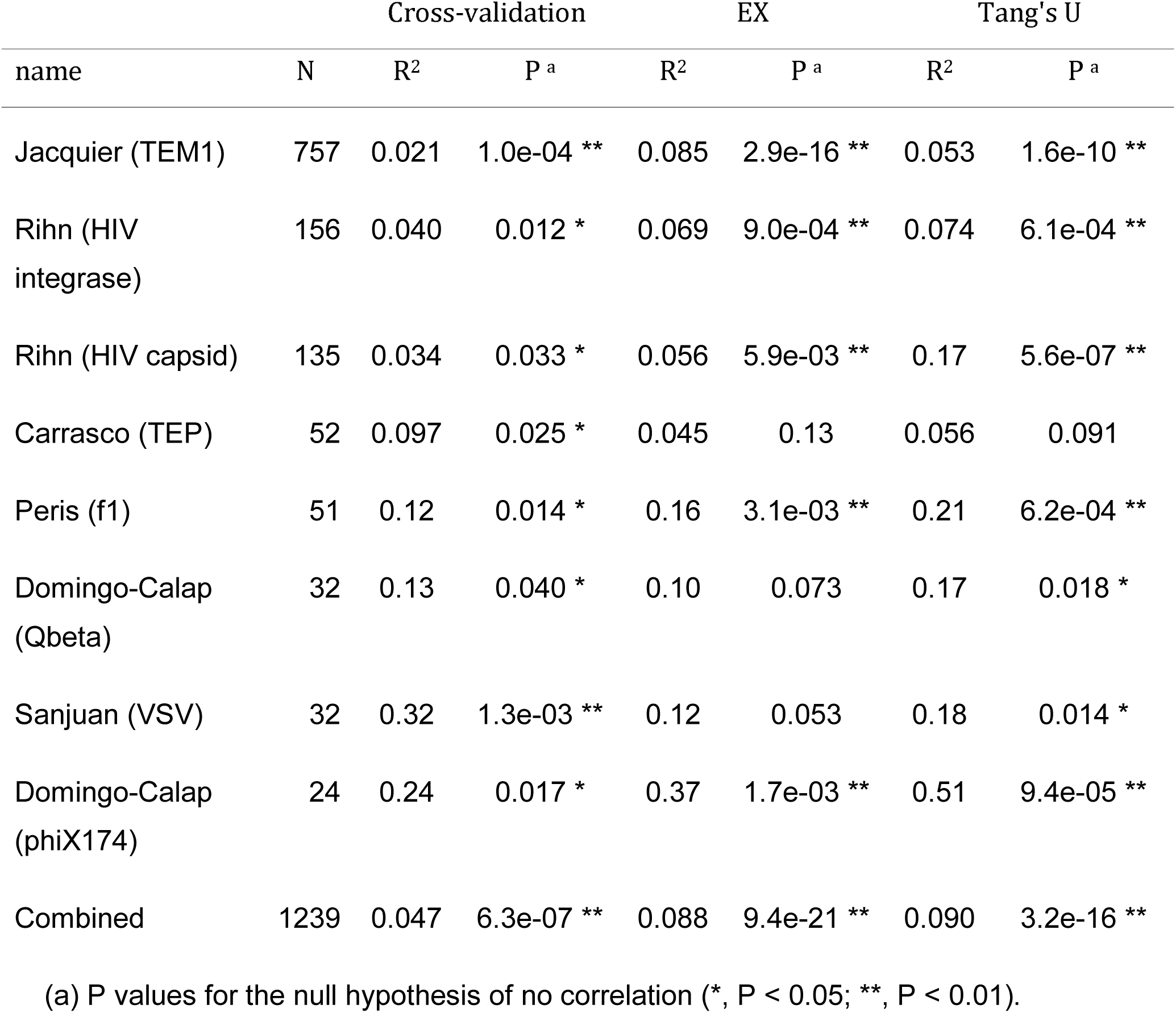
Power of 8 studies to discern generic effects of exchangeability.

| name | N | Cross-validation |  | EX |  | Tang's U |  |
| --- | --- | --- | --- | --- | --- | --- | --- |
|  |  | R<sup>2</sup> | P<sup>a</sup> | R<sup>2</sup> | P<sup>a</sup> | R<sup>2</sup> | P<sup>a</sup> |
| Jacquier (TEM1) | 757 | 0.021 | 1.0e-04 ** | 0.085 | 2.9e-16 ** | 0.053 | 1.6e-10 ** |
| Rihn (HIV integrase) | 156 | 0.040 | 0.012 * | 0.069 | 9.0e-04 ** | 0.074 | 6.1e-04 ** |
| Rihn (HIV capsid) | 135 | 0.034 | 0.033 * | 0.056 | 5.9e-03 ** | 0.17 | 5.6e-07 ** |
| Carrasco (TEP) | 52 | 0.097 | 0.025 * | 0.045 | 0.13 | 0.056 | 0.091 |
| Peris (f1) | 51 | 0.12 | 0.014 * | 0.16 | 3.1e-03 ** | 0.21 | 6.2e-04 ** |
| Domingo-Calap (Qbeta) | 32 | 0.13 | 0.040 * | 0.10 | 0.073 | 0.17 | 0.018 * |
| Sanjuan (VSV) | 32 | 0.32 | 1.3e-03 ** | 0.12 | 0.053 | 0.18 | 0.014 * |
| Domingo-Calap (phiX174) | 24 | 0.24 | 0.017 * | 0.37 | 1.7e-03 ** | 0.51 | 9.4e-05 ** |
| Combined | 1239 | 0.047 | 6.3e-07 ** | 0.088 | 9.4e-21 ** | 0.090 | 3.2e-16 ** |

(a) P values for the null hypothesis of no correlation (\*, P < 0.05; \*\*, P < 0.01).

### Correlation of mutant fitnesses with amino acid exchangeability

To assess the power of mutation studies individually and collectively, we correlate observed mutant fitness quantiles with expected values from three independent predictors: the EX matrix (Yampolsky and Stoltzfus 2005), the U matrix of Tang, et al (2004), and a cross-validation predictor. The cross-validation predictor applied to a given target study is constructed from all **other** studies (i.e., excluding the target study), and is simply a matrix of mean quantiles for each type of replacement (e.g., Ala to Val).

EX and U are used on the grounds of being powerful and mutationally unbiased predictors, whereas various biochemical predictors are less powerful (as will become apparent below), and various evolution-based measures other than U (e.g., PAM, BLOSUM), though perhaps powerful, cannot be used, because they are not known to be free of the mutational effects that we wish to exclude. The EX matrix, based on a meta-analysis of early mutation studies (which reported phenotypes other than fitness), was designed specifically to serve as a mutationally unbiased measure of exchangeability in models that separate selection from mutation. In a comparative evaluation, EX was shown to be as powerful, or more powerful, than a representative sample of other predictors (Yampolsky and Stoltzfus 2005). The “universal evolutionary index” or U matrix of Tang, et al (2004) is based on modeling evolution of thousands of genes, using a method designed to separate codon-level mutational effects from protein-level effects. It purports to be a measure of evolutionary acceptability that scales directly with the rate of evolution.

The results of using EX, U and a cross-validation predictor (Table 1) indicate that even small studies of mutant fitnesses have considerable power to reveal generic effects of amino acid exchangeability. For instance, for the study of 135 HIV capsid mutants by Rihn, et al., there is a significant correlation between the fitness reported for a mutant and the predictor for the relevant replacement type (e.g., Val to Ala), whether the predictor is EX, U, or a cross-validation predictor based on the other studies. This shows, not only that individual studies are powerful, but that there is a consistency across studies: although most effects of an amino acid replacement in a protein are very context-dependent (which is why the R^2^ values are small), generic effects of exchangeability are seen across sites and proteins.

### Ability to distinguish conservative from radical replacements

The conservative transitions hypothesis proposes that transitions collectively are more conservative than transversions. How well do mutant fitness studies distinguish conservative replacements from radical ones? We construct two versions of this distinction, EX_B_ and U_B_ (the “B” indicates a binary distinction, as opposed to a continuous measure), simply by designating higher-exchangeability replacements as “conservative”, and the remainder as “radical”.

Table 2 shows how well studies of mutant fitness distinguish conservative from radical replacements, and how well they distinguish transitions from transversions. The measure of effect-size used here is the chance that a mutant designated as “conservative” is more fit than a randomly chosen “radical” mutant. This statistic is not affected by the relative sizes of the 2 classes; its range is from 0 to 1, with a null expectation of 0.5; higher values indicate that nominally “conservative” changes are indeed conservative. We call this measure AUC because it has the same meaning as the area under a ROC (receiver-operating characteristic) curve for a binary classifier. That is, as pointed out by Hanley and McNeil (1982), the AUC for a binary classifier is equivalent to the chance that a randomly chosen positive instance will be ranked higher than a randomly chosen negative instance (see Methods).

**Table 2.** Power of 8 studies to detect various binary distinctions.

| name | N | EX<sub>B</sub> |  | U<sub>B</sub> |  | TiTv |  |
| --- | --- | --- | --- | --- | --- | --- | --- |
|  |  | AUC | P<sup>a</sup> | AUC | P<sup>a</sup> | AUC | P<sup>a</sup> |
| Jacquier (TEM1) | 757 | 0.66 | 2.8e-14 ** | 0.61 | 4.0e-08 ** | 0.54 | 0.019 * |
| Rihn (HIV<br>integrase) | 156 | 0.64 | 2.1e-03 ** | 0.67 | 7.2e-05 ** | 0.540 | 0.191 |
| Rihn (HIV capsid) | 135 | 0.60 | 7.1e-03 ** | 0.63 | 7.4e-04 ** | 0.50 | 0.50 |
| Carrasco (TEP) | 52 | 0.64 | 0.037 * | 0.61 | 0.087 | 0.50 | 0.50 |
| Peris (f1) | 51 | 0.60 | 0.13 | 0.72 | 4.6e-03 ** | 0.54 | 0.31 |
| Domingo-Calap<br>(Qbeta) | 32 | 0.60 | 0.16 | 0.65 | 0.076 | 0.75 | 0.084 |
| Sanjuan (VSV) | 32 | 0.61 | 0.13 | 0.75 | 6.1e-03 ** | 0.31 | 0.93 |
| Domingo-Calap<br>(phiX174) | 24 | 0.81 | 5.8e-03 ** | 0.94 | 1.6e-04 ** | 0.34 | 0.85 |
| Combined | 1239 | 0.65 | 6.5e-18 ** | 0.64 | 1.6e-12 ** | 0.53 | 0.024 * |

(a) P values for a one-sided test where the alternative is that nominally conservative replacements, or transitions, are more fit (\*, $P < 0.05$ ; \*\*, $P < 0.01$ )

Even small studies have significant power to distinguish conservative from radical substitutions based on EX_B_ and U_B_. In the combined data set, the AUC is 0.65 for EX_B_ and 0.64 for U_B_. That is, a conservative replacement according to EX_B_ has a 65 % chance of being more fit than a randomly drawn radical replacement.

However, the same studies typically do not distinguish transitions from transversions. Only one study shows a marginally significant result (p = 0.019 for the largest study). The combined results for the entire set of 1239 replacements are shown at the bottom of Table 2. For the combined data, the AUC is 0.53, with a 95 % confidence interval of 0.50 to 0.56 (based on 400 bootstrap replicates).

One might object that this approach is framed incorrectly, in that it uses the entire distribution of mutational effects, whereas the distribution of changes fixed in evolution is obviously weighted toward more modest effects, because natural selection removes the most damaging ones whether they are transitions or transversions. If the changes actually accepted in evolution are mostly in the top 50 %, or the top 5 %, of the fitness distribution, then this is the fraction that should be examined most closely to test the conservative transitions hypothesis.

The effect of testing for a ti:tv effect at successively higher thresholds of fitness is shown in Figure 1. In fact, the AUC does not go up if we filter out the low end, but stays close to 0.5.

**Figure 1.**
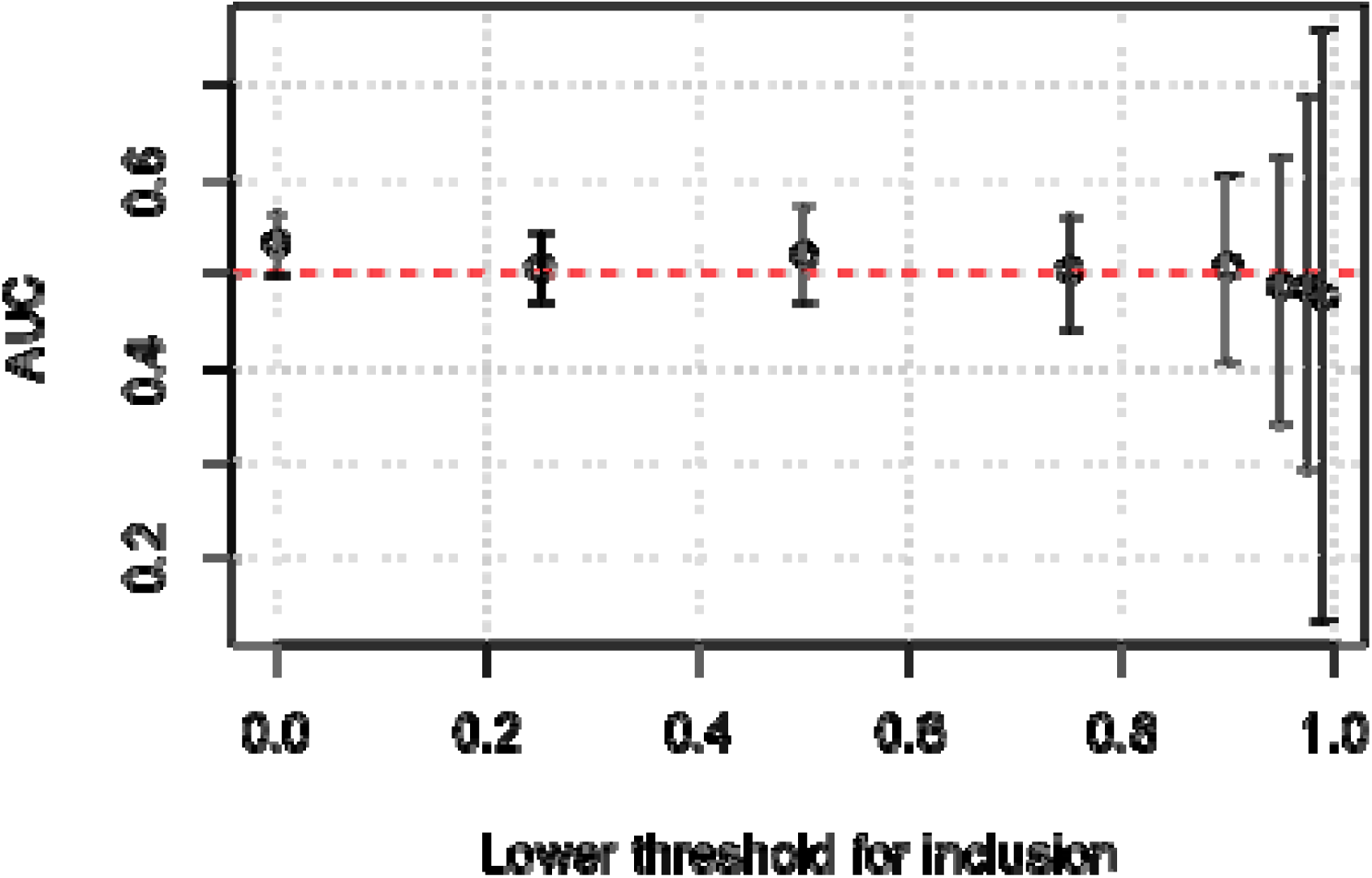
The relative conservativeness of transitions for distributions of mutant effects truncated at the low end of fitness. The advantage of transitions (AUC) is shown as a function of threshold quantile for left-truncated data, e.g., the AUC value for x = 0.2 is computed without the bottom 20 % of the distribution. Under the conservative transitions hypothesis, one might expect that, even if there is no advantage over the entire distribution, an advantage will appear at the high end. In fact, this is not observed. As mentioned in the text, AUC = 0.53 for the complete set of data, corresponding to a truncation threshold of 0, i.e., no truncation. As the threshold increases, AUC decreases (rather than increases), though the differences are insignificant.

Another way to explore the upper end of the fitness distribution is to consider studies of mutational effects that focus on **beneficial** mutations (Ferris, et al. 2007; MacLean, et al. 2010; Miller, et al. 2011; Schenk, et al. 2012). These studies are small, with only 15 to 38 mutants, and have little power to distinguish effects of exchangeability. For the combined set of 111 beneficial mutants shown in Table 3, the AUC for the conservative transitions hypothesis is 0.40 (95 % CI, 0.28 to 0.51), suggesting that perhaps beneficial transitions are not more, but less fit than beneficial transversions.

**Table 3.** Relative advantage of transitions in 4 studies of beneficial mutations

| name | N | EX<sub>B</sub> |  | U<sub>B</sub> |  | TiTv |  |
| --- | --- | --- | --- | --- | --- | --- | --- |
|  |  | AUC | P<sup>a</sup> | AUC | P<sup>a</sup> | AUC | P<sup>a</sup> |
| Schenk (TEM1) | 38 | 0.46 | 0.63 | 0.40 | 0.84 | 0.49 | 0.53 |
| MacLean (RpoB) | 31 | 0.45 | 0.68 | 0.47 | 0.62 | 0.35 | 0.93 |
| Miller (ID11) | 27 | 0.58 | 0.30 | 0.62 | 0.18 | 0.25 | 0.96 |
| Ferris (phi6) | 15 | 0.69 | 0.18 | 0.64 | 0.20 | 0.39 | 0.76 |
| NA | 111 | 0.52 | 0.50 | 0.50 | 0.57 | 0.40 | 0.95 |

(a) P values for a one-sided test where the alternative is that nominally conservative replacements, or transitions, are more fit (\*, $P < 0.05$ ; \*\*, $P < 0.01$ )

### Gauging the evolutionary effect size

As mentioned above, the conservative transitions hypothesis has 2 parts, a claim that transitions are conservative, and a claim that this conservativeness accounts for an evolutionary pattern. The present set of studies suggests that transitions are more conservative, but only slightly. How important could an effect of this size be?

One way to ask this question is to compare the ti:tv distinction to various biochemical distinctions. Any quantitative property of an amino acid can be used to create a conservative-vs-radical distinction, e.g., for a measure of the polarity of each amino acid, the “conservative” changes will be the ones with the least change in polarity. The AAindex database (Kawashima and Kanehisa 2000) has data on nearly 250 biochemical factors (see Methods). The random sample of 25 factors from AAIndex shown in Figure 2 indicates that biochemical predictors typically are (1) considerably more powerful than the ti:tv distinction, and (2) considerably less powerful than EX and U (to view the full set of predictors, see Supplementary Material, section 3).

**Figure 2.**
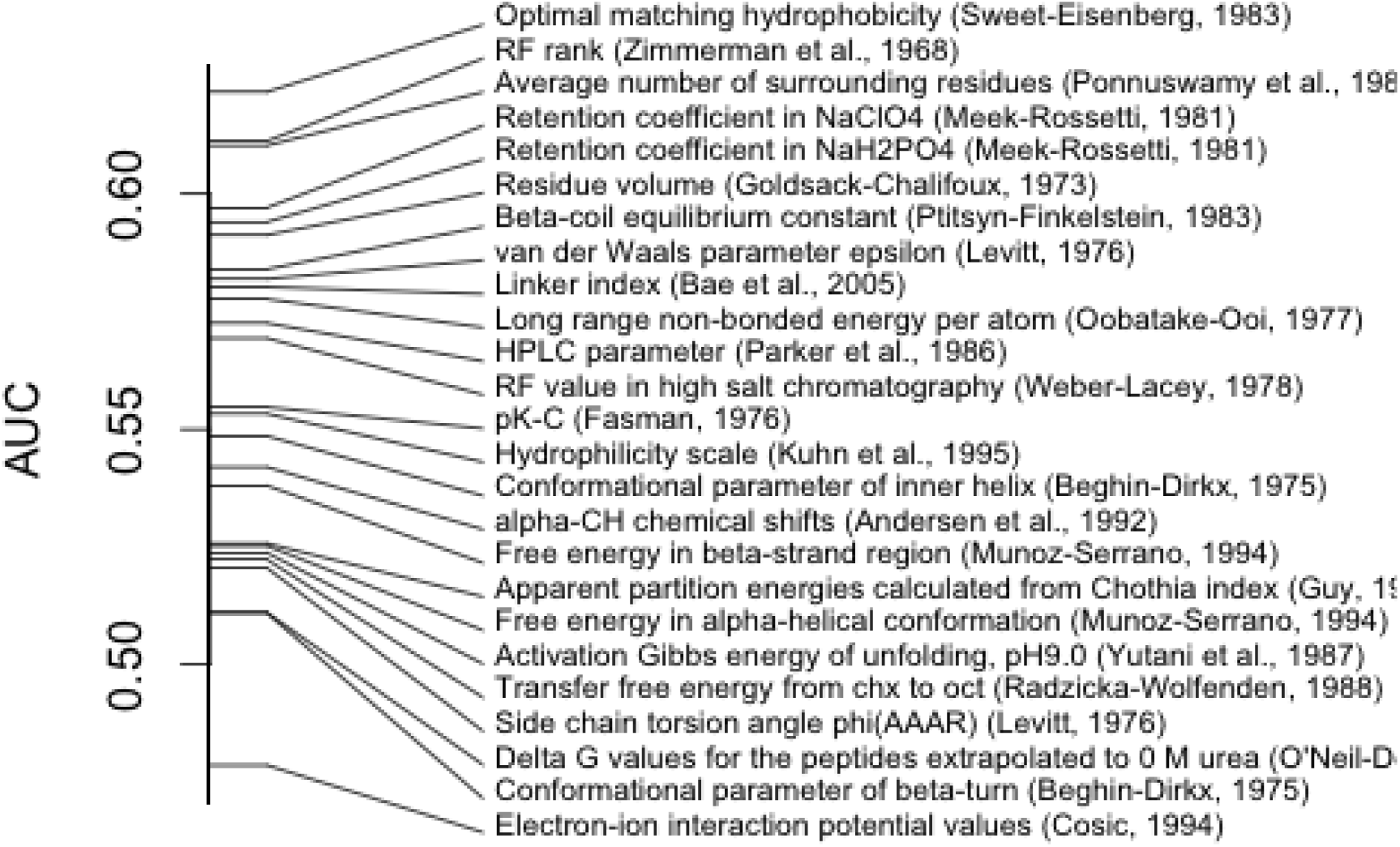
Relative power of conservative-radical distinctions based on some biochemical factors. Binary predictors based on 25 randomly chosen biochemical factors from the AAIndex database were applied to the prediction of mutant fitnesses in mutation-scanning experiments (for the full set of predictors, see Supplementary Material, section 3). The AUC is the chance that a randomly chosen mutant designated as “conservative” by the predictor has a higher fitness than one designated as “radical”. Most predictors are more powerful than the ti:tv distinction (AUC = 0.53).

Yet, natural selection has the ability to amplify small differences into major effects. Perhaps a difference with an effect size of AUC = 0.53 might translate into a several-fold bias in terms of evolutionary acceptance.

How do these two relate to each other? The U matrix illustrates this relationship, because values of U scale with evolutionary rates, and U_B_ has a known power as a conservative-radical distinction, namely AUC = 0.64. The ratio of U values for conservative replacements relative to radical ones is 2.7. That is, conservative replacements as defined by U_B_ are 2.7-fold more likely to be accepted in evolution than radical ones.

This pair of values, AUC = 0.64 and evolutionary bias = 2.7, represents one point in the relationship between evolutionary acceptability and classification power for mutant fitness effects. There is another point where AUC = 0.5 (no power) and evolutionary bias = 1 (no effect). We can fill in the relationship further by randomizing U_B_, as shown in Figure 3. The results show that, when about 75 % of the values are randomized, U_B_ has an AUC of 0.53, equal to that of the transition:transversion distinction. This corresponds to an evolutionary bias of 1.3. The confidence interval of AUC from 0.50 to 0.56 for the transition:transversion distinction corresponds to the interval of 1.0 to 1.6 in evolutionary bias. That is, the expected evolutionary effect of the transition:transverion bias is a 1.3-fold bias, with a confidence interval from 1.0 (no effect) to 1.6. This makes it unlikely that selection plays the major role in causing the evolutionary transition:transversion bias, which typically is several-fold favoring transitions.

**Figure 3.**
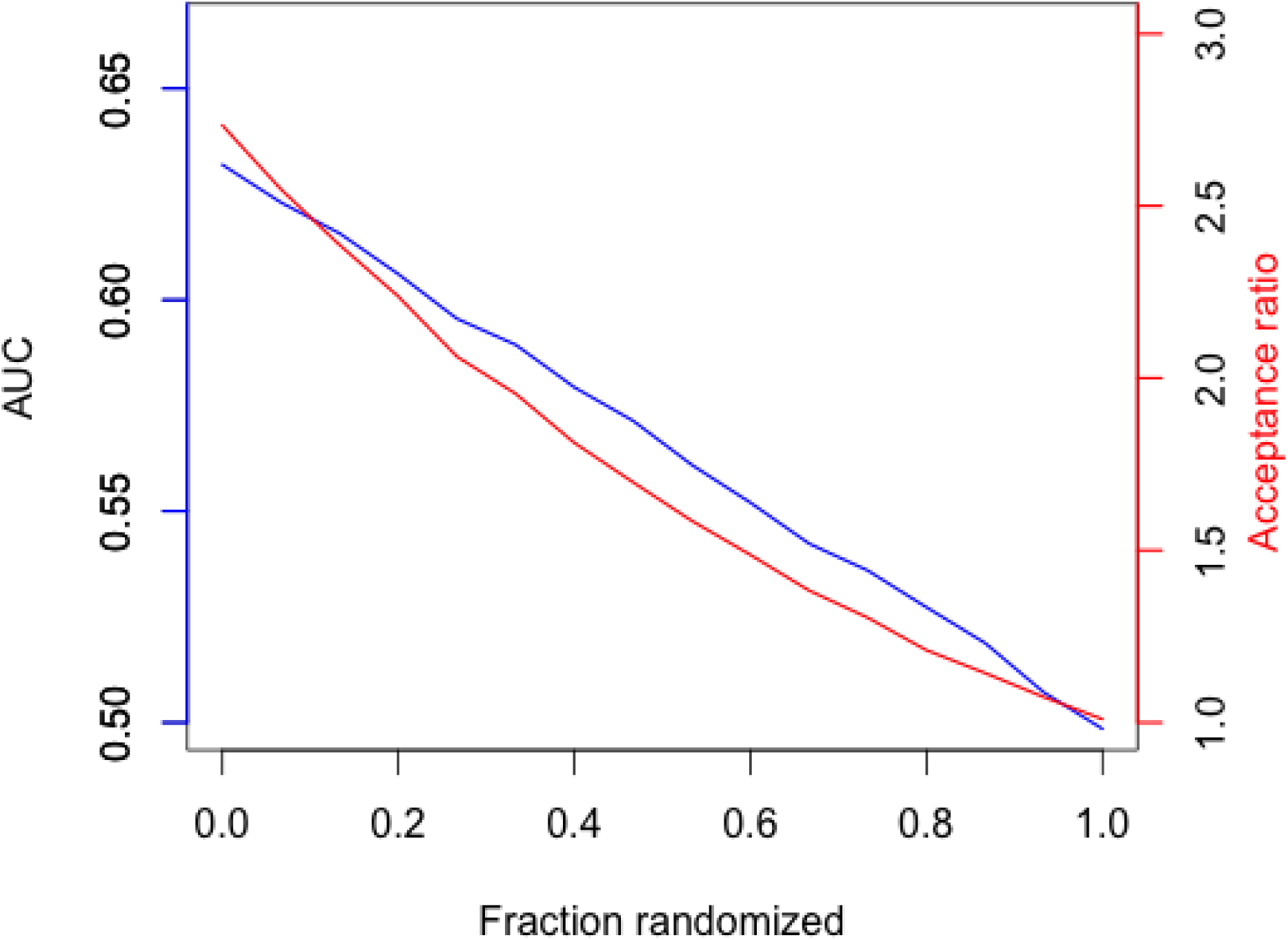
Relationship between power to predict mutant fitnesses and evolutionary effect size. AUC and evolutionary acceptance ratio are shown for increasingly randomized verions of U_B_. For the unrandomized U_B_, the power in predicting mutational al effects is AUC = 0.64, and this corresponds to an evolutionary acceptance ratio of 2.7 for conservative versus radical replacements. To estimate the evolutionary acceptance ratio for more modest values of AUC, we can weaken U_B_ by randomly re-assigning “conservative” or “radical” labels to an increasingly large fraction of replacement types (200 replicates at each level of randomization). The AUC of 0.53 is reached at about 75 75 % randomization, where the evolutionary effect size is 1.3.

### Discussion

Based on a collection of 8 studies that report fitnesses for replacement mutations, we have assessed the prospects for the hypothesis that the conservativeness of replacements via transition accounts for their increased frequency in evolution. Even small studies reveal predictable patterns of amino acid exchangeability, and most have sufficient power to distinguish a binary conservative-vs-radical distinction. However, the same studies typically do not show significant conservativeness of transitions. Overall, the chance of a transition mutation being more fit than a transversion is 53 % (95 % CI, 50 % to 56 %). This effect size is not large compared to that of most biochemical predictors, and is not large enough to explain the several-fold bias toward transition replacements observed in evolutionary studies.

The finding that the conservativeness of transitions is a rather weak effect increases the prospects for the alternative mutational explanation, in which the rate at which new alleles are introduced by transition mutations is several-fold higher than for transversions, and this bias predisposes evolutionary change to happen via transitions.

Though this idea may be familiar, it relates to a rather substantial and unresolved issue in evolutionary genetics, which is the extent to which evolution in nature happens in the “gene pool” regime imagined by the architects of the Modern Synthesis, in the kind of mutation-driven regime imagined by early mutationists and later molecular evolutionists, or something in between (see McCandlish and Stoltzfus 2014). For many, years the mutationist view has suffered from an association with neutral evolution (e.g., the two remain conflated in Nei 2013). This may explain the ongoing popularity of the conservative transitions hypothesis: a mutation bias is the obvious explanation for transition bias in the evolution of introns and other non-coding sequences, but was not accepted for protein sequences, which are assumed to be “under selection” and thus not subject to such biases. Yet it is clear theoretically that mutation-biased adaptation is possible and, more generally, that mutation and selection can both contribute to orientation or direction in evolution (Yampolsky and Stoltzfus 2001; Stoltzfus 2006).

The results presented here also prompt the question of how the lore that transitions are conservative was estabished. In a survey of the literature, we found that, when the alleged conservativeness of transitions is attributed to a source, the source is often Zhang (2000), or early works such as Fitch (1967), Grantham (1974) or Vogel and Kopun (1977). Grantham does not directly address this issue, but a genetic-code-based calculation shows that the mean Grantham distance for transition-mediated replacements is lower than that for transversions, e.g., as indicated in Table 2 of Xia, et al (1998). The study by Vogel and Kopun is often cited as evidence for the conservative transitions hypothesis, because they present a calculation that, for 3 different biochemical measures, suggests that transitions are more conservative. Nevertheless, Vogel and Kopun themselves favored a mutational explanation (see hypothesis 3 on p. 179), arguing that the effect size is too small for conservativeness of transitions to account for the evolutionary bias.

These prior studies are inconclusive for 2 very general reasons. The first is that none reports an effect size sufficient to account for the evolutionary bias. For instance, Zhang's (2000) analysis of 3 possible conservative:radical distinctions finds that the distinction based on Miyata, et al (1979) yields the largest evolutionary effect size, which is a 2-fold effect, i.e., radical replacements are roughly half as likely to accumulate, relative to null expectations. However, though the effect of conservativeness is strong, the link reported between transitions and conservativeness is weak. According to Zhang (2000), the chance that a transition is conservative by Miyata's measure is 35 %, compared to 33 % for transversions, a proportional difference of only 6 % (i.e., 2 / 33 = 0.06). Miyata-conservativeness may be a 2-fold evolutionary effect, but if transitions are only 6 % more Miyata-conservative than transversions, the overall bias will be far less than 2-fold.

Second, none of these works escapes the kind of logical circularity pointed out by di Giulio (2001; see also Yampolsky and Stoltzfus 2005), in which a measure of evolutionary tendencies (e.g., PAM, BLOSUM) is invoked to argue for effects of selection rather than mutation, ignoring the fact that the pattern of evolution is influenced (to an unknown degree) by mutational effects. This is an indirect form of the Panglossian fallacy, i.e., it is formally a fallacy of arguing that transitions are more adaptive simply because they happen more often, without inquiring into why they happen more often.

The fallacy is not avoided by invoking biochemical factors. The popular composite indices of “biochemical” distance constructed by Grantham (1974) and Miyata, et al (1979) are based on choosing biochemical factors that fit well with observed evolutionary patterns from earlier protein comparisons. Likewise, all 3 “biochemical” measures used by Vogel and Kopun (1977) are based on fitting to protein comparisons. The problem with this approach is suggested by Figure 4, which shows the conservativeness of transitions for a random sample of the biochemical indices in the AAindex database (Kawashima and Kanehisa 2000). Roughly half make transitions seem conservative, and the other half make them seem radical.

**Figure 4.**
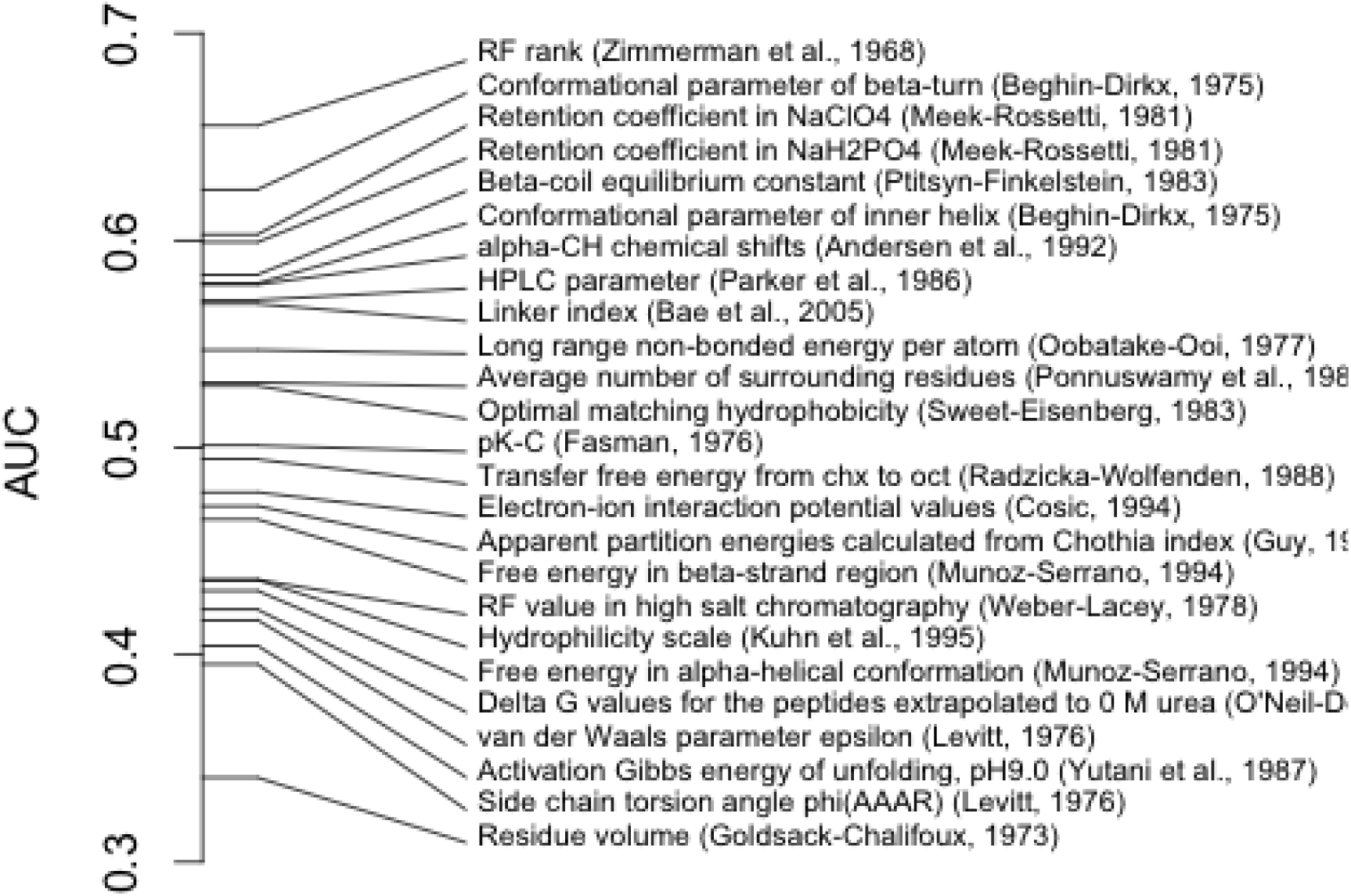
The relative advantage of transitions as indicated by a random sample of biochemical factors. Each biochemical attribute of an amino acid is converted to a pairwise similarity measure, so that each possible amino acid replacement has a similarity score. Here the AUC is the chance that a replacement due to a randomly chosen transition (from the pool of actual mutants from the 8 studies) has a higher similarity score (for the given biochemical attribute) than a randomly chosen transversion. The resulting distribution indicates that many biochemical factors make transitions seem more conservative (AUC > 0.5), but a roughly equal number make transversions seem more conservative (AUC < 0.5). For the full set of predictors, see Supplementary Material, section 3.

As shown earlier (Figure 2), this is not because biochemical indices are generally poor predictors of exchangeability. Instead, among many moderately powerful predictors, there are ones that make transitions seem favorable, and others that make transversions seem favorable. Thus, converting evolutionary patterns into biochemical descriptors before re-applying them to the analysis of evolutionary patterns does not allow one to escape a logical circularity: if early molecular evolutionists had found that transversions dominate evolution, they could have rationalized this pattern by appeal to biochemistry just as easily as they rationalized the observed dominance of transitions.

## Materials and Methods

### Identification of studies and data sets for inclusion

An initial core set of studies (Sanjuan, et al. 2004; Carrasco, et al. 2007; Roscoe, et al. 2013) was expanded by including other work cited by these studies. Then this set was expanded further by open-ended searches based on keywords or by tracking citations. In general, no text-based search does a good job of recovering mutation-scanning studies of the desired type. Narrow searches (e.g., “distribution of mutational effects”) implicate only a fraction of true positives and did little to expand the core set of studies; broad searches (e.g., “mutation” plus “fitness”) implicate so many false positives that they are impractical and were abandoned. Most relevant studies cite the pioneering work of Sanjuan (2004) or the seminal review by Eyre-Walker and Keightley (2007). Candidate studies identified in this manner were screened for appropriateness, ultimately resulting in the 8 studies listed in Table 1. The search covered literature published through December, 2014 and does not include more recent studies.

As noted in Materials and Methods, we restricted our attention to studies with (1) a size of at least 20 replacement mutants; (2) measures of growth (fitness) rather than simply biochemical activities; and (3) a random or arbitrary set of mutants. Most excluded studies of mutational effects have only a few mutants, or they report effects on binding or activity (but not on fitness), or they are focused on achieving particular outcomes rather than exploring a random set of variants, or they use deep sequencing to identify and quantify mutants, an approach that introduces uncontrolled nucleotide biases (Supplementary Material, section 2).

### Processing and management of mutation data

Starting from raw data tables supplied by authors (either directly, or via published supplements), all further processing and analysis steps were encoded in scripts. For each study used here, there is a an R-Markdown (Rmd) file that (when executed in an appropriate environment, such as RStudio) describes and executes the steps (e.g., cleaning, re-coding, sequence integration) to convert input data into a standard tabular form in which there is a single row describing each mutant and its effects. The figures and tables in this paper are generated by further Rmd scripts that operate on the standardized input data.

### Other data sources

Values of U are from Tang, et al. (2004), and values of EX are from published supplements. Biochemical indices from the AAIndex database were accessed via the Interpol package (Heider 2012) and custom R code.

Note that, although AAindex lists 533 biochemical indices, less than half are pure biochemical indices. The others are based on some method of counting occurrences in naturally evolved proteins, e.g., frequency with which an amino is found in a helix. Because the distribution of an amino acid in an evolving set of natural proteins will depend on the distributions of its closest mutational neighbors, such measures are not mutationally unbiased. They were removed using a custom list of name exclusion patterns (“[fF]requenc”, “[pP]reference”, “[cC]ompositi”, “[pP]ropensit”, “[dD]istribution”, “[iI]nformation”, “[wW]eights”, “[oO]ccurrence”, “[Pp]roportion”, “probability”, “mutability”, “Geisow”, “Janin”), resulting in a set of 247 indices.

### Tests of power and effect

The results presented here rely mainly on standard statistical procedures. When P values are reported in Table 1 for a linear predictor, this is from the t-test in the built-in linear model (lm) function in R. When P values are reported for binary predictors in Tables 2 and 3, this is based on the Wilcoxon-Mann-Whitney test as implemented in the “wilcox.test” function of the R “stats” package, using a one-sided test. When confidence intervals are given on an AUC value, this is based on re-sampling using 400 bootstrap replicates.

The only unfamiliar methods involve the use of binary predictors. To convert a biochemical index C to a binary distinction, we first convert it to a pairwise similarity by the formula Sij = 1 − abs(C_i_ − C_j_) / max, where max is the maximum absolute difference. Converting a continuous measure of similarity into a binary measure is a simple matter of assigning all values above a particular quantile to the “conservative” class, and the rest to the “radical” class. To ensure that a constructed predictor is comparable to the transition:transversion distinction, the threshold is chosen so that the conservative class is the same size as the transition class in the data to be tested.

As explained above, we can define a measure of effect-size with intuitive properties that we designate as AUC, based on an application of ROC analysis that may not be obvious. In ROC analysis of a binary classifier, each instance has a binary state (e.g., disease vs. non-disease), and the classifier makes a ranking of instances and predicts the binary state based on a threshold. The ROC curve plots the true-positive rate against the false-negative rate, and the area under this curve is equivalent to the chance that a randomly chosen positive instance is ranked higher than a randomly chosen negative one (Hanley and McNeil 1982). If we treat the fitness study as the classifier that supplies a ranking for each mutant, and the conservative-radical distinction as the binary state of a mutant, then the AUC is the chance that a mutant of a nominally conservative type has a higher fitness than a randomly chosen mutant of a nominally radical type. The relationship of AUC to the Wilcoxon-Mann-Whitney test is explained by Manley (1982). Calculating AUC from the test statistic is an algebraic conversion based on the formula AUC = (pairs – WMW_statistic(x, y)) / pairs, where x and y are vectors representing the two samples, and pairs = length(x) * length(y). This formula applies specifically to wilcox.test in the R “stats” package (some other implementations define the test statistic in a different way).

Note that converting fitnesses to within-study quantiles allows us to compare studies, and allows us to combine data for across-study tests. The use of quantiles rather than absolute fitnesses does not have any effect on a within-study AUC or Wilcoxon-Mann-Whitney test, which is non-parametric.

## Acknowledgements

We thank Greg Babbitt and David McCandlish for comments, and Ashley Acevedo, Rafael Sanjuan and Nicholas Wu for help in acquiring and interpreting data. The identification of any specific commercial products is for the purpose of specifying a protocol, and does not imply a recommendation or endorsement by the National Institute of Standards and Technology.

